# MPKaDB: A p*K*_a_ Database for Exploring pH Dependence in Membrane Proteins

**DOI:** 10.1101/2025.11.14.688495

**Authors:** Jiahao He, Yansheng Chen, Jinxi Wu, Zhitao Cai, Wenting Jia, Qiuchen Yue, Yandong Huang

## Abstract

The biological activities of many membrane proteins are pH-regulated, yet mapping their pH dependence experimentally is slow and expensive. In this work, we present MPKaDB (http://computbiophys.com/DeepKa/mpkadb), a comprehensive p*K*_a_ database for membrane proteins that instantly decode the protonation states of ionizable residues under a specified pH. Leveraging MPKaDB, we performed pH-coupled electrostatic characterization of trans-membrane proteins. To facilitate use, a user-friendly search engine was developed to retrieve a protein of interest and returns its p*K*_a_ values, isoelectric points of both cytoplasmic and extra-cytoplasmic faces, and an automated screening of active-site residues. In the end, two case studies were proposed to demonstrate how p*K*_a_’s from MPKaDB could be applied to explore the pH-dependent relationship between membrane protein structure and function.

**Graphical Abstract:** 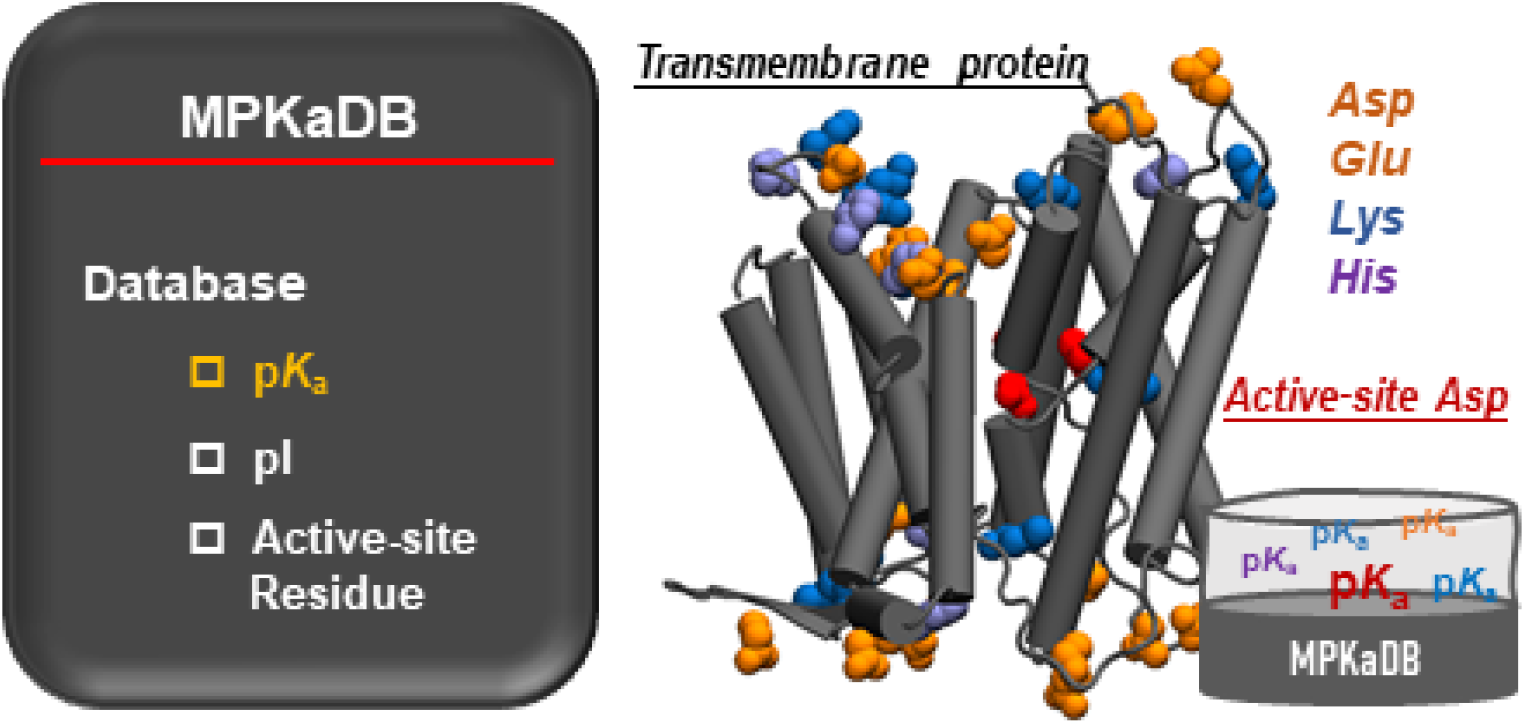

## 1. Introduction

pH plays a central role in many biological processes, including transports of ions or small molecules across the cellular membrane by membrane-embedded proteins, such as sodium-proton antiporters [1], proton channels [2], glucose-proton symporters [3], and cholesterol transporters [4]. Normally, such a proton-coupled mass transport event occurs under a specific pH window, exhibiting high pH dependence. For instance, the secondary sodium-proton antiporter NhaA of *Escherichia coli* functions only when the pH is higher than 6.5 and the reactivity reaches the maximal around pH 8.5 [5]. Thus, to understand the underlying mechanism, it is essential to identify the proton carriers according to the p*K*_a_ values of ionizable residues in the reaction core [6]. Above sodium transports are driven by the pH gradient maintained primarily by proton channels, activities of which are governed by pH too [2]. Beyond screening the candidates of proton carriers, an accurate description of the salt-bridge network lining the proton-conducting path is equally critical, and this also hinges on reliable p*K*_a_ calculations of ionizable residues that involve in salt bridging [7].

More than ten thousand crystal structures of transmembrane proteins have been experimentally determined. Based on the average occurrence (Fig. S1A) and transmembrane-centre likelihood (Fig S1B) of each ionisable residue type determined from about one thousand non-redundant transmembrane proteins in the OPM database [8], ionizable residues are essential constituents of reaction cores typically situated in the middle of the membrane and their titration behaviors determined by p*K*_a_’s should be considered explicitly, especially those carried out near physiological pH. In addition, there is a considerable high probability of encountering an acid and a base simultaneously (FIg. S1C), implying the importance of coupling between ionizable residues.

Evidently, p*K*_a_ values are indispensable for linking protein function to pH. However, experimental p*K*_a_ values for membrane proteins remain scarce, probably due to pH-triggered conformational changes during the transport cycle. For example, it was found that the inward-facing gate of NhaA opens in response to the deprotonation of an essential Asp residue [6], providing a direct connection between side-chain protonation state and large-scale conformational switching. Thus, accurate p*K*_a_ predictions in theory become urgent. Constant pH molecular dynamics (CpHMD) simulations have been demonstrated applicable to transmembrane proteins [6]. In particular, CpHMD simulations explicitly couple changes in protonation state to conformational rearrangements, enabling direct estimation of macroscopic p*K*_a_’s within a lipid-bilayer environment. However, CpHMD simulations are time demanding and therefore impractical for the high-throughput p*K*_a_ screens demanded by industrial work flows.

Alternatively, DeepKa [9], a deep learning-based scheme, has been applied to probing the influence of conformational changes sampled by MD simulations on p*K*_a_’s and the resulting deprotonation equilibria [10, 11]. Notably, ionizable residues in the active sites are often deeply buried within the proteins and therefore hidden from lipids. As a result, the environment that an ionizable side chain senses would still be dominated by the protein, which to some extent explains the feasibility of DeepKa working on trans-membrane proteins. Recently, performances of several fast p*K*_a_ predictors, including the artificial intelligence (AI)-based DeepKa [12], the empirical formula Propka [13] and Poisson-Boltzmann (PB) equation-based H++ [14], on the outer membrane porin F (OmpF) from *Escherichia coli*, a trimeric channel, was evaluated using CpHMD as benchmark [15]. It was found that DeepKa outperforms the widely used PropKa and H++, in consistence with the benchmark by Wei et al. on p*K*_a_’s of ionizable residues that belong to water-soluble proteins [16].

In this work, the p*K*_a_ database MPKaDB that contains p*K*_a_ data of 3.3 million Asp, Glu, His and Lys residues in 12 k membrane proteins deposited in the protein data bank (PDB) was created by DeepKa [9]. The database is freely downloadable from the web page of MPKaDB (http://computbiophys.com/DeepKa/mpkadb) or GitLab (https://gitlab.com/computational-biophysics/deepkaserver/-/tree/main/DeepKaDB). Additionally, a user-friendly interface is available for searching membrane proteins of special interest via PDB codes. Notably, the MPKaDB web page has been integrated into the existing DeepKa web server [10]. In the result section, electrostatics of transmembrane proteins were characterized based on charges determined by p*K*_a_ values and pH. Besides, two case studies were set out to demonstrate how the p*K*_a_ database could be applied to preliminarily exploring the pH-governed mechanisms of transmembrane proteins.

## 2. Materials and Methods

### 2.1. Data Collection, Annotation and Access

The present MPKaDB contains p*K*_a_ values estimated by DeepKa for 3,306,058 Asp, Glu, His, and Lys residues in 12,602 membrane proteins from the PDB. Cd-hit [17] combined with BLAST+ [18] was utilized to screen non-homogeneous proteins from eight thousand transmembrane proteins in OPM. The sequence identity threshold of 30 percent was applied. As a result, a cluster of 968 non-homogeneous transmembrane proteins was obtained to characterize titration-associated electrostatics of transmembrane proteins.

All p*K*_a_’s of MPKaDB were saved in a CSV file that contains five columns where the first four together locate a residue and the last is the p*K*_a_ value. The CSV file can be downloaded freely from the MPKaDB page of the DeepKa web server or our GitLab page. Apart from the one-click downloading of the entire database. The MPKaDB page also contains a search engine. To search a membrane protein of interest, users are supposed to enter a valid PDB code. If the membrane protein is not available, users are recommended to use the Prediction page where both PDB codes and files are acceptable inputs [10].

### 2.2 Isoelectric Point (pI) Calculation

pI is the pH value at which a molecule carries zero net charge. Charges of Asp/Glu/Cys/Tyr (*q*^a^) and His/Lys/Arg (*q*^b^) residues that vary as a function of pH were calculated by Eq. 1 and Eq. 2, respectively, equivalent to the Henderson-Hasselbalch (HH) formula [19]. Here the reference p*K*_a_’s measured in aqueous solvent, 8.55, 9.84 and 12.0, were utilized for Tyr, Cys and Arg, respectively, for pI calculations [20]. On the other hand, to compute the p*K*_a_ shifting in response to protein environment, reference p*K*_a_ values of 3.67, 4.25, 6.54 and 10.4 were applied to Asp, Glu, His and Lys, respectively [20]. In addition to pI, such pH-dependent charge calculations will be utilized to the membrane-associated statistics of electrostatics.

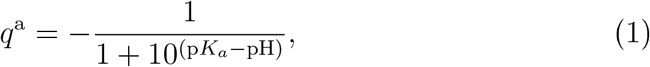

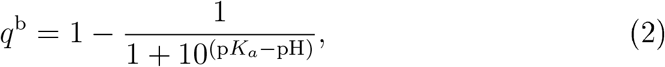

### 2.3. Molecular Dynamics (MD) Simulation

All-atom MD simulations of a glucose-proton symporter (GlcP_*Se*_, PDB code: 4LDS) [3], a proton pump (NsXeR, PDB code 6EYU) [21] and an anion channel (iC++, PDB code 6CSM) [22] were set up to investigate the coupling between protonation state and electrostatics in pores. Details about the system preparation and simulation protocols can be found in the Supporting Information. Finally, protein structures were visualized by VMD [23] and analyzed by MDAnalysis [24].

## 3. Results and Discussion

### 3.1. MPKaDB Web Server Description

The web server provides two core functions, a one-click button to download the entire MPKaDB database as a ZIP-compressed archive, and a search engine that retrieves any entry via a valid PDB code (Fig. 1A). If the protein of interest is found, a clickable arrow directs the user to its Result page that contains two columns. As shown in Fig. 1B, the left column includes a link to download the PDB file. Below are the pI^Cyto^ (z≤0) and pI^Extr^ (z>0) that correspond to cytoplasmic (Cyto) and extra-cytoplasmic (Extr) domains, respectively, and the total charges varying as the pH are visualized.

**Figure 1:**
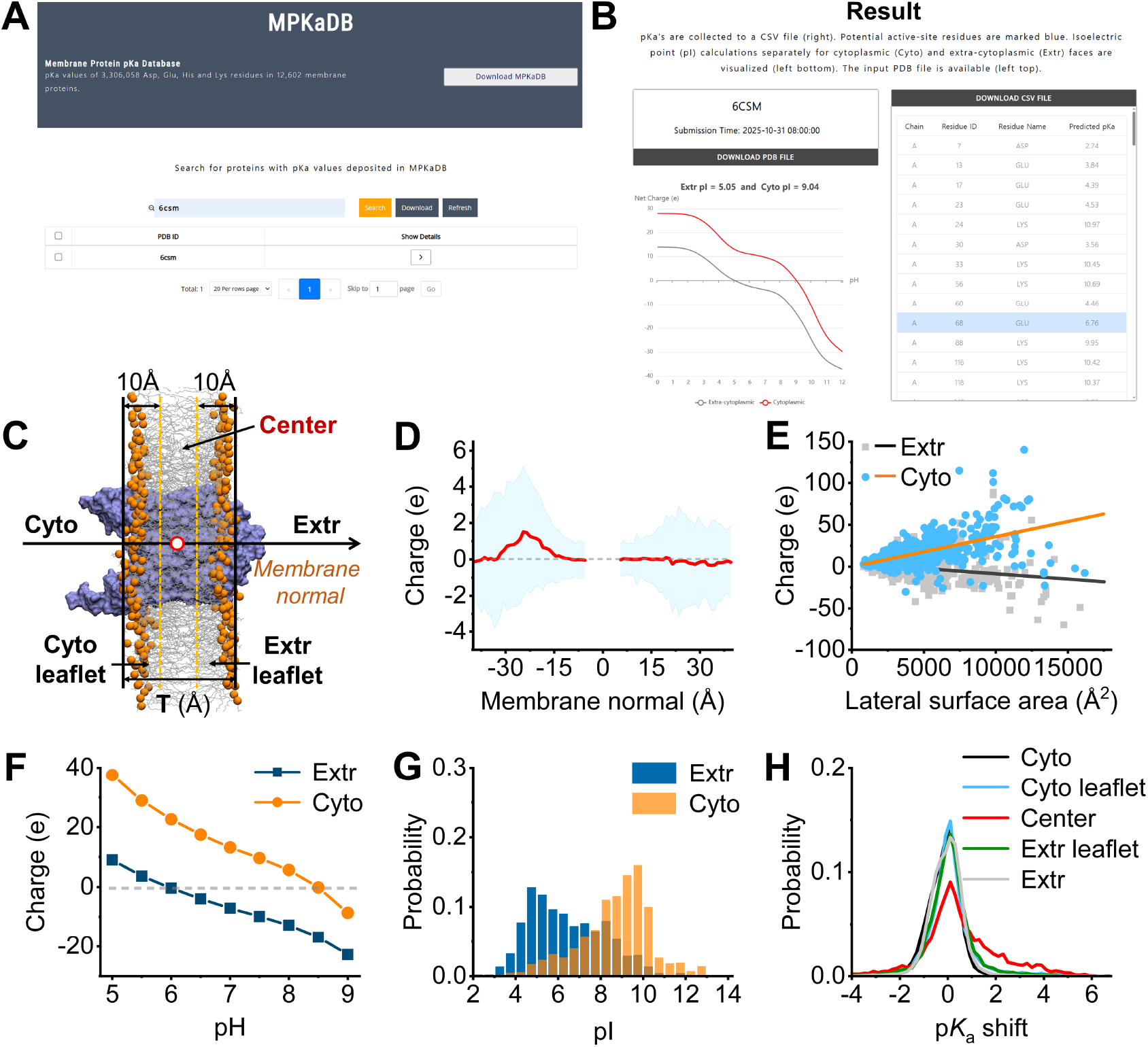
MPKaDB web page and downstream analyses. MPKaDB (A) front and (B) result page. (C) Region definitions across the membrane. The protein is displayed with the surface mode. Lipids are displayed with lines and P atoms at lipid heads are highlighted with orange spheres. The origin was marked by a open red circle. (D) Mean (red) and standard deviation (cyan) of net charge as a function of z coordinate. (E) Correlation plots between lateral surface areas ( Å^1^) and mean net charges as well as regression fitting lines for Extr (gray and black) and Cyto (blue and orange), respectively. The correlation coefficients are 0.54 and −0.24 for Cyto and Extr, respectively. (F) Mean net charge as a function of pH at Extr (blue) and Cyto (orange), respectively. (G) pI distributions at Extr (blue) and Cyto (orange), respectively. (H) p*K*_a_ shift distributions at five regions, Cyto (black), Cyto leaflet (blue), Center (red), Extr leaflet (green) and Extr (gray), respectively, across the membrane.

The right column contains the other link to download the p*K*_a_ data as a CSV le. As one can see, p*K*_a_ values are enumerated on the web page as well. Residues highlighted in blue are screened by the active-site residue criteria that the p*K*_a_ value ranges from 5 to 9 and simultaneously the z coordinate of the C*α* atom is within the central region of the membrane (Center) as shown in Fig. 1C. Specifically, for each transmembrane protein in the OPM database, the membrane thickness (T, around 30 Å) was used to define the outer limits of the Cyto and Extr leaflets, −T/2 and T/2, respectively. The distance from the outer boundary of either leaflet towards the Center was set to 10 Å. As a result, the scope of Center is T-20 Å.

### 3.2. Electrostatic Characterization of Transmembrane proteins

The positive-inside rule, a fundamental phenotype of membrane asymmetry [25], for transmembrane proteins has long been an consensus [26, 28] and is critical to membrane-protein topology [27]. Previous studies, however, simply assumed Asp/Glu and Lys/Arg to be negatively and positively charged, respectively, without estimating pH-associated charges. Now that p*K*_a_ values in MPKaDB enable the charge of an ionizable residue to vary as a function of pH according to the HH equation, in this work electrostatic properties across the membrane were revisited based on the screened set of non-homogeneous transmembrane proteins.

As shown in Fig. 1D, the peak positive net charge at the physiological pH seven is −24 Å, about 10 Å away from the count maximal of Arg or Lys nearby the Cyto leaflet [26, 28]. Interestingly, negative net charge, though weak, at the distance of 15 Å from the Extr leaflet is observed, consistent with the statistics by Baker et al. [28]. The biological relevance of the positions for positive and negative net charges in average at two sides of the membrane are not clear. Given protein structures, the correlation between protein lateral surface area and the net charge in ranges of −T/2*±*10 for Cyto and T/2*±*10 for Extr. As illustrated in Fig. 1E, the absolute net charge scales linearly with the lateral surface area on both the Cyto and Extr, revealing, albeit modest, an interplay between molecular size and net charge.

Furthermore, the mean net charges at different pH conditions were computed and shown in Fig. 1F, from which one can tell that the averaged pI values for Extr and Cyto are 6.0 and 8.5, respectively. Here we assume that charges at Cyto could be fully isolated from those at Extr by the low-dielectric membrane, which leads to two pI values, one for Cyto and the other for Extr. pI values for individual proteins were computed too. As illustrated in Fig. 1G, The pI distribution for Cyto is more concentrated when compared with that for Extr, exhibiting high certainty of the positive charge preference at the cytoplasmic side.

p*K*_a_’s are the basis of aforementioned analyses. To examine the p*K*_a_ shifting in response to the membrane, distributions of p*K*_a_ deviations from reference p*K*_a_’s, namely p*K*_a_ shifts, were computed in five regions over the membrane normal. From Fig. 1H, it is evident that large p*K*_a_ shifts are more populated while approaching the Center region. For one thing, an ionizable residue prefers the neutral state when buried, which would explain large p*K*_a_ shifts. For another, the likelihood (Fig. S1) of finding only Asp/Glu (23.2%) in the Center region is much higher than that for Lys/Arg (9.1%), which would further explain the bias of p*K*_a_ shifting up for a buried Asp/Glu residue. Such a membrane-coupled p*K*_a_ shifting highlights the necessity of taking titration behaviors into account while investigating electrostatic properties of transmembrane proteins.

### 3.3. Case Study I: Identification of Proton Carriers in Sodium-Proton Antiporters

Sodium-proton antiporters are well known by pH-regulated translocation of ions. Structures of electrogenic NhaA in *Escherichia coli* and NapA from *Thermus thermophilushave* that share the same fold type, characterized by two anti-parallel discontinuous helices crossing over in core domains, have been extensively studied. As collected in Tab. 3.3, eight and three crystal structures are available for NhaA and NapA, respectively. Although the experimental pH scans from the inactive 4.0 to the optimum 8.5, all existing crystal structures of NhaA should correspond the resting inward-facing (IF) state, judging from radius of gyrations for both IF and outward-facing (OF) gates (Tab. S1). In contrast, either the IF or OF structure was obtained under the optimal pH of 8.0 for NapA.

Either NhaA or NapA transports one sodium by exchanging two protons. However, the identity of the two residues that carry the protons remains disputed. To address this question, p*K*_a_’s of ionizable residues in the active site for these structures were extracted from MPKaDB. Taking NhaA as an example, Asp163, Asp164, and Lys300 buried in the reaction center (Tab. S1) have been proposed as candidate proton carriers. Asp164 as one proton carrier has been a consensus, thus the controversy focuses on Asp163 and Lys300. In general, p*K*_a_’s for Asp163 and Asp164 are higher than those reported from CpHMD simulations, most notably for Asp163, which likely reflects insufficient conformational sampling of the allosteric motions unique to NhaA [6, 10, 11]. Notably, we found that p*K*_a_ of Asp164 is correlated to the solvent accessible surface area (SASA) of the carboxyl oxygens on its side chain (Fig. S2), which implies that the titration of Asp164 could be dominated by hydrophobic forces. As to Lys300, all p*K*_a_’s (Tab. 3.3) cluster at or just above the reference value of 10.4, except for the 1ZCD structure determined at the inactive pH of 4.0, where the loss of the Asp163–Lys300 salt bridge (Fig. S3A) shifts the p*K*_a_ downward (Fig. S3B). A more thorough discussion of this divergence can be found in two previous studies [6, 10].

Regarding Asp133 also buried in the reaction center, we found that p*K*_a_ values are all lower than 5.0, well below the functional pH optimum of 8.5 for NhaA (8.0 for NapA), in line with previous CpHMD data on NhaA [6], except for structure 7A0X, which was determined at pH 6.0 (Tab. 3.3). Asp133 side chain has been suggested to precisely neutralize the positive charge that accumulates between two helical termini (Fig. S4A) [29]. We note that Asp133 is also surrounded by three hydrophobic residues, namely Lue18, Phe136 and Phe339, which would induce p*K*_a_ to shift up (Fig. S4B), allowing Asp133 to carry a proton, supporting Asp133 as an alternative proton-binding site proposed by Shen group [36]. Thus, p*K*_a_ values curated in MPKaDB support Asp163, Asp164 and Lys300 in NhaA as viable proton-carrier candidates and Asp133 remains a plausible backup.

p*K*_a_’s of counter-residues in NapA, namely Glu333, Asp156, Asp157 and Lys305, that correspond to Asp133, Asp163, Asp164 and Lys300, respectively, are also provided. Notably, the p*K*_a_ values of the three Asp residues are modestly higher in OF, in line with NhaA [11], whereas that of Lys300 is slightly lower, compared with IF. Finally, based on the mean p*K*_a_ values (Tab. 3.3), the same ensemble of candidate proton carriers identified for NhaA could be applied to NapA as well.

**Table 1:** p*K*_a_’s of Na^+^/H^+^ antiporters NhaA and NapA deposited in PDB.

| PDB<br>ensemble | pH | State | Active site $pK_a$'s | Ref |
| --- | --- | --- | --- | --- |
| NhaA | 8.5 |  | D133/D163/D164/K300 |  |
| 1ZCD | 4.0 | IF | 4.3/7.8/7.0/8.2 | [29] |
| 4AU5 | 4.0 | IF | 4.7/5.7/6.0/11.3 | [30] |
| 4ATV<sup>a</sup> | 4.0 | IF | 4.6/6.6/5.5/11.0 | [30] |
| 7A0X | 6.0 | IF | 5.7/7.6/8.2/11.3 | [31] |
| 7S24 | 6.5 | IF | 4.0/7.7/7.8/11.5 | [32] |
| 8PS0 | 7.5 | IF | 4.4/6.5/6.0/10.7 | [33] |
| 7A0Y<sup>b</sup> | 8.2 | IF | 4.8/6.6/6.8/none | [31] |
| 7A0W | 8.5 | IF | 4.7/6.4/6.5/11.0 | [31] |
| Mean |  |  | 4.6/6.9/6.7/10.7 |  |
| NapA | 8.0 |  | E333/D156/D157/K305 |  |
| 4BWZ | 7.8 | OF | 4.0/6.6/5.7/10.9 | [34] |
| 5BZ2 | 8.0 | IF | 4.5/7.2/6.9/10.3 | [35] |
| 5BZ3 | 8.0 | OF | 3.4/5.7/6.0/10.8 | [35] |
| Mean |  |  | 4.0/6.5/6.2/10.7 |  |

<sup>a</sup> Triple mutants of A109T, Q277G, and L296M.
<sup>b</sup> Single mutant of K300R.

### 3.4. Case Study II: Coupling between Protonation State and Salt Bridging in Transmembrane Proteins

In this case study, p*K*_a_’s (Tab. S2) combined with MD simulations were employed to probe how protonation states modulate the salt-bridge networks that line mass-transport pathways. Here three transmembrane proteins named GlcP_*Se*_ [3], NsXeR [21] and iC++ [22] were selected based on the active-site residue criteria above. As illustrated in Fig. 2, residues that satisfy the criteria above are Asp22 in GlcP_*Se*_, Asp76 in NsXeR and Glu68/Asp234 in iC++. Notably, the calculated p*K*_a_ of Asp22 (Tab. S pka of three acids) coincides with the pH optimum 5.5, identifying this residue as the most probable H^+^-binding site of the proton pump GlcP_*Se*_ [3].

**Figure 2:**
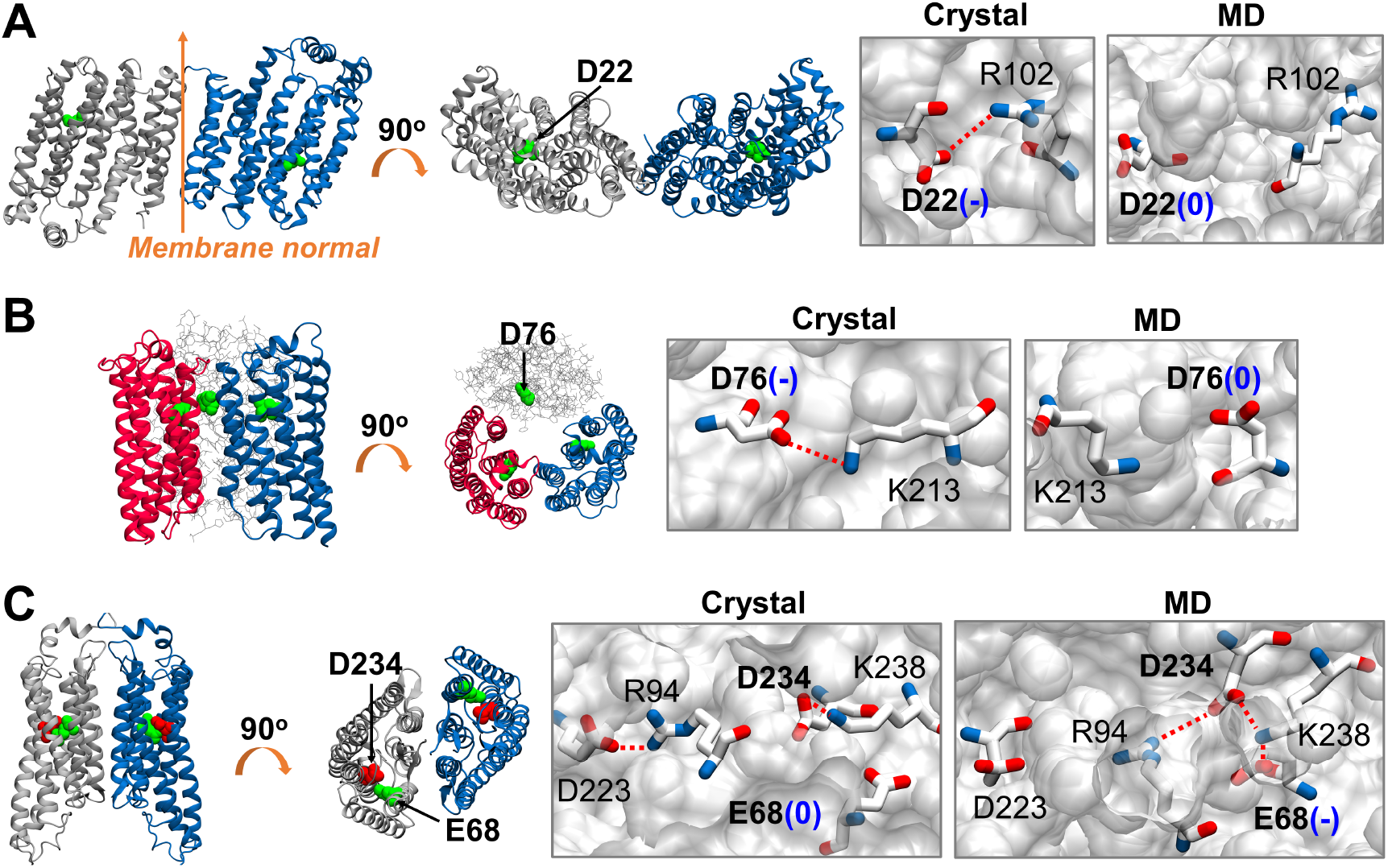
Crystal structures and salt bridge networks of three transmembrane proteins. Crystal structures are visualized with the new cartoon model and chains A, B and C are colored gray, blue and red, respectively. Acids screened are displayed with green or red spheres. Residues involved in salt bridge networks are displayed with sticks where carbon, oxygen and nitrogen atoms are colored white, red and blue, respectively. Side and top views of crystal structures and the comparisons of salt bridge networks in pores between crystallized and MD simulated structures for proteins (A) GlcP_*Se*_, (B) NsXeR and (C) iC++, respectively. The orange arrow represents the membrane normal. Dashed red lines represent salt bridges. Symbols - and 0 in parentheses indicate deprotonated and protonated states of Asp or Glu, respectively.

Judging from the crystal structures, Asp22 of GlcP_*Se*_ (Fig. 2A) and Asp76 of NsXeR (Fig. 2B) are in contact with the positively charged Arg102 and Lys213, respectively, and therefore are likely to be charged or deprotonated (Tab. S3). In contrast, Glu68 of iC++ remains too far from Lys238 to form a salt bridge (Fig. 2C), and thus would take the neutral or protonated state (Tab. S3). Noting that these initial guesses were subsequently validated by fixed-charge MD simulations (Fig. S5-10), it is of interest to see what would happen if the protonation states adopted by crystal structures were changed.

As illustrated in Fig. 2A, neutralization of Asp22 completely breaks the Asp22-Arg102 salt bridge (Fig. S5) and swings the Arg102 side chain toward the tunnel entrance. On the other hand, as shown in Fig. 2B, the Asp76-Lys213 salt bridge is disrupted when Asp76 is neutralized (Fig. S6). Unlike GlcP_*Se*_, Lys213 does not turn away and would still interact with Asp76.

The salt-bridge networks above are simple when compared with that found in iC++. As illustrated in Fig. 2C, when Glu68 is charged, it forms a stable salt bridge with Lys238 (Fig. S7), which nevertheless does not affect the Lys238-Asp234 salt bridge (Fig. S8). Interestingly, the Arg94-Asp223 salt bridge nearby the entrance is disrupted (Fig. S9). Arg94 turns to forms a new salt bridge with Asp234 (Fig. S10), which affects the stability of the Lys238-Asp234 salt bridge (Fig. S11).

Intriguingly, though Glu68 is distant from Asp223, the Asp223-Arg94 salt bridge is governed by the titration of Glu68. Such a long-distance coupling between ionizable residues was also observed in NhaA by Henderson et al. [36]. When Glu68 is charged, the net charge at the Center becomes negative, which might explain the attraction of Asp94 towards the Center to neutralize the charge and therefore preserve electrostatic equilibrium.

## 4. Conclusions and Perspectives

Here we present MPKaDB, a p*K*_a_ database dedicated exclusively to membrane proteins. The current release catalogs 3.3 million Asp, Glu, His and Lys residues from 12,006 membrane protein structures deposited in the PDB. The complete database is available for free download as a single compressed archive from the MPKaDB website. A user-friendly protein search engine has also been implemented where users can instantly retrieve any entry by entering its PDB code. Beyond p*K*_a_ values, the result page delivers downstream annotations, including an initial screening of potential active-site residues and the isoelectric points calculated separately for the cytoplasmic and extra-cytoplasmic faces. Finally, MPKaDB will be updated in parallel with the PDB repository.

Equipped with p*K*_a_ data, we dissected the electrostatic properties of membrane proteins, including the well-known positive-inside rule, with pH-resolved detail. Two case studies further illustrate how the database could be used, in conjunction with pH, to probe membrane protein structure–function relationships. In the first case study, by integrating PDB structures with *in silico* screening for potential active-site residues, we evaluate the likelihood of individual residues serving as proton carriers in two sodium – proton antiporters. Whereas, in the second case study, MD simulations were used to probe how changing protonation states modulate the salt-bridge networks at protein centers. Notably, we revealed a long-distance coupling between two acidic residues, which is hardly captured by a static PDB structure.

The current MPKaDB omits Cys, Tyr, and Arg because DeepKa intrinsically cannot predict their p*K*_a_ values, asking for the development of a general-purpose model in the future. However experimental or theoretical p*K*_a_ data for the three reidue types are extremely scarce [37], which poses a major challenge for model development in future. Besides, DeepKa was trained on water-soluble proteins. Thus, residues located nearby the water-lipid interface should be warranted, as DeepKa treats the lipid phase as water, which would underestimate p*K*_a_ shifts. Finally, pI values at cytoplasmic and extra-cytoplasmic faces were computed separately, which asks for precedent spatial arrangements of membrane proteins with respect to the hydrocarbon core of the lipid bilayer from the third-part resources, like OPM and PDBTM [38]. Thus, the pI field on the Result page will remain empty if the protein is unavailable in either OPM or PDBTM.

## 6. CRediT authorship contribution statement

Yandong Huang: Writing, Supervision, Project administration, Conceptualization, Validation, Methodology. Jiahao He, Yansheng Chen, Jinxi Wu: Software, Resources, Methodology. Zhitao Cai: Validation, Methodology. Wenting Jia and Qiuchen Yue: Software.

## 7. DATA AVAILABILITY

The database is freely available as a web server at http://computbiophys.com/DeepKa/mpkadb or GitLab at https://gitlab.com/computational-biophysics/deepkaserver/-/tree/main/DeepKaDB.

## Appendix A. Supplemental data

Supplementary material to this article can be found online.

