## Supporting Information for "MPKaDB: A p*K*_a_ Database for Exploring pH Dependence in Membrane Proteins"

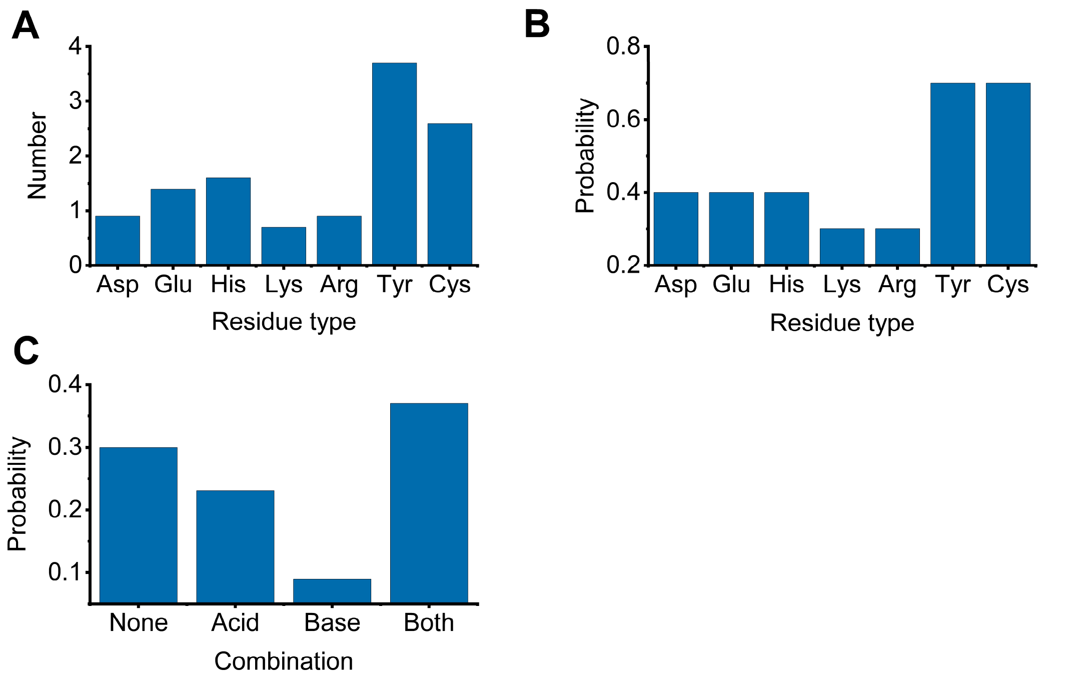


**Figure S1.** (A) Mean numbers of seven ionizable residues in the middle of the membrane. (B) the likelihood of finding an ionizable residue type in the middle of the membrane. (C) Probabilities of combinations of acids (Asp, Glu) and bases (Lys, Arg) in the middle of the membrane. None indicates neither acids nor bases are present. Acid/Base indicates at least one acid/base is present and bases/acids are absent. Both indicates at least one acid and one base are present. Statistics are based on 968 non-homogeneous transmembrane proteins.

**Table S1.** Radius of gyration (Rg) of inward-facing (IF) and outward-facing (OF) gates as well as solvent accessible surface areas (SASA) of four ionizable residues in the reaction core of NhaA.

| PDB code | IF gate Rg (Å)[a] | OF gate Rg (Å)[b] | SASA (Å$^2$)[c] | | | |
|---|---|---|---|---|---|---|
| | | | Asp133 | Asp163 | Asp164 | Lys300 |
| 1ZCD | 4.2 | 4.5 | 0 | 3.4 | 0 | 0 |
| 4AU5 | 4.5 | 4.7 | 0 | 3.0 | 3.7 | 0 |
| 4ATV | 4.2 | 4.7 | 0 | 14.7 | 14.9 | 0 |
| 7A0X | 4.4 | 4.6 | 3.9 | 6.8 | 1.8 | 0 |
| 7S24 | 4.4 | 4.6 | 1.6 | 2.7 | 0 | 0 |
| 8PS0 | 4.4 | 4.8 | 8.5 | 11.7 | 7.9 | 0 |
| 7A0Y | 4.4 | 4.6 | 0 | 5.1 | 2.6 | 0 |
| 7A0W | 4.3 | 4.7 | 7.2 | 10.0 | 3.4 | 0 |

[a] IF gate is composed by sidechains of Val75, Ile34, Met157, Ala160 and Ile161[1].

[b] OF gate is composed by sidechains of Phe72, Ala167, Ile168, Phe344 and Ile345[2].

[c] SASA is computed by LCPO/NLR model[3, 4] reparaemterized with CAHRMM force field [5].

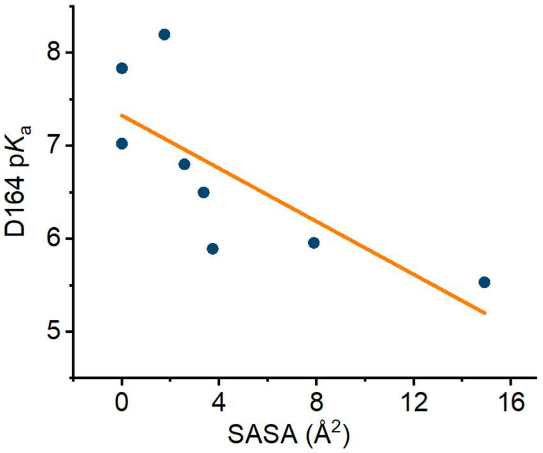


**Figure S2.** Correlation plot between SASA and p$K_a$ values for Asp164 in the eight PDB structures of NhaA. The Pearson's correlation coefficient is -0.75.

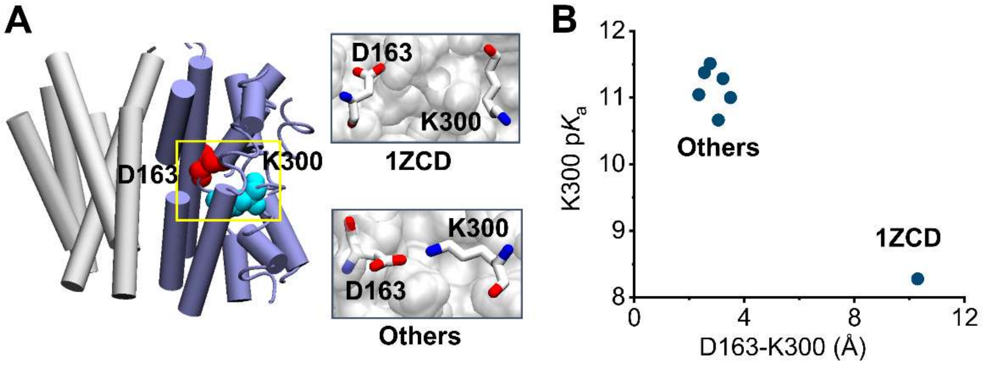


**Figure S3.** (A) Structure (PDB code: 1ZCD) of NhaA displayed with the cartoon model. Scaffold and core domains are colored gray and ice blue, respectively. Asp163 and

Lys300 in the yellow frame are shown with red and cyan spheres. The interaction between Asp163 and Lys300 for the PDB structure 1ZCD (up) and others (down) that represent other seven PDB structures of NhaA are visualized with sticks. (B) Correlation plot between the minimal distance between Asp163 and Lys300 side chains and p$K_a$ values of Lys300 based on eight structures of NhaA.

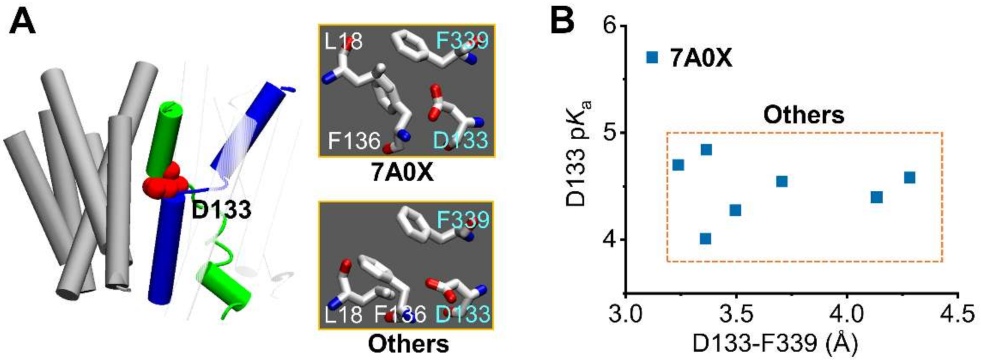


**Figure S4.** Structure of NhaA (PDB code: 4A5U) displayed with the cartoon model. Scaffold domain is colored gray. Two crossed discontinuous helices IV and XI at core domain are colored green and blue, respectively. Asp133 is displayed with red spheres. Lue18, Asp133, Phe136 and Phe339 for the PDB structure 7A0X (up) and Others that represent other seven PDB structures of NhaA (down) are shown with sticks. (B) Correlation plot between the minimal distance between the minimal distance between Asp133 carboxyl oxygens and Phe339 benzene ring and p$K_a$ values of Asp133 based on eight structures of NhaA.

**Table S2.** p$K_a$ values of ionizable residues identified as potential active-site residues.

| PDB code | Residue | p$K_a$ | | |
|---|---|---|---|---|
| | | Chain A | Chain B | Chain C |
| 4LDS | Asp22 | 5.4 | 5.3 | n/d |
| 6EYU | Asp76 | 6.0 | 4.3 | 5.9 |
| 6CSM | Glu68 | 6.8 | 5.3 | n/d |
| | Asp234 | 5.2 | 4.3 | n/d |

**Table S3.** Minimal distances between Asp/Glu side chain oxygens and Lys/Arg side chains nitrogens.

| PDB code | Residue pair | Distance (Å) | | |
|---|---|---|---|---|
| | | Chain A | Chain B | Chain C |
| 4LDS | D22-R102 | 4.7 | 5.0 | n/d |
| 6EYU | D76-K213 | 4.4 | 3.9 | 4.2 |
| 6CSM | E68-K238 | 5.4 | 5.8 | n/d |
| | D234-E238 | 3.5 | 3.5 | n/d |

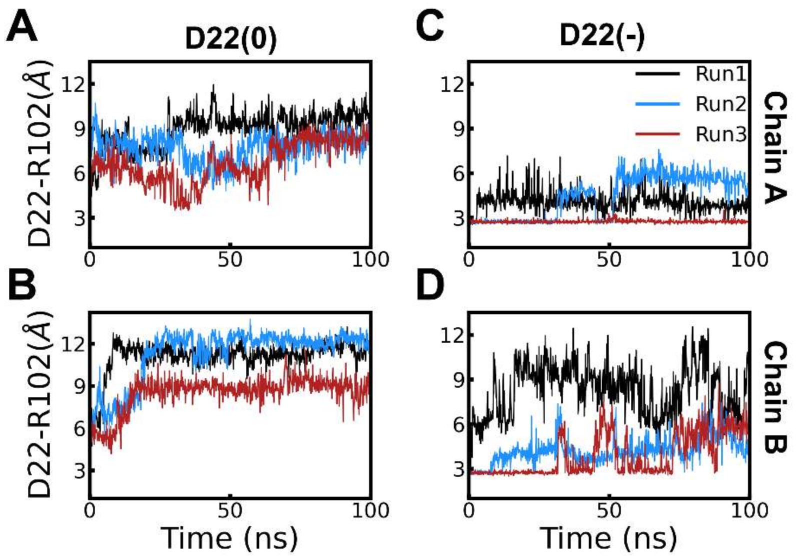


**Figure S5.** MD simulation trajectories of the minimal distance between Asp22 and Arg102 side chains in Glucose-proton symporter GlcP$_{Se}$ where Asp22 is fixed in (A-B) protonated and (C-D) deprotonated states, respectively. Simulations started from the crystal structure (PDB code: 4LDS).

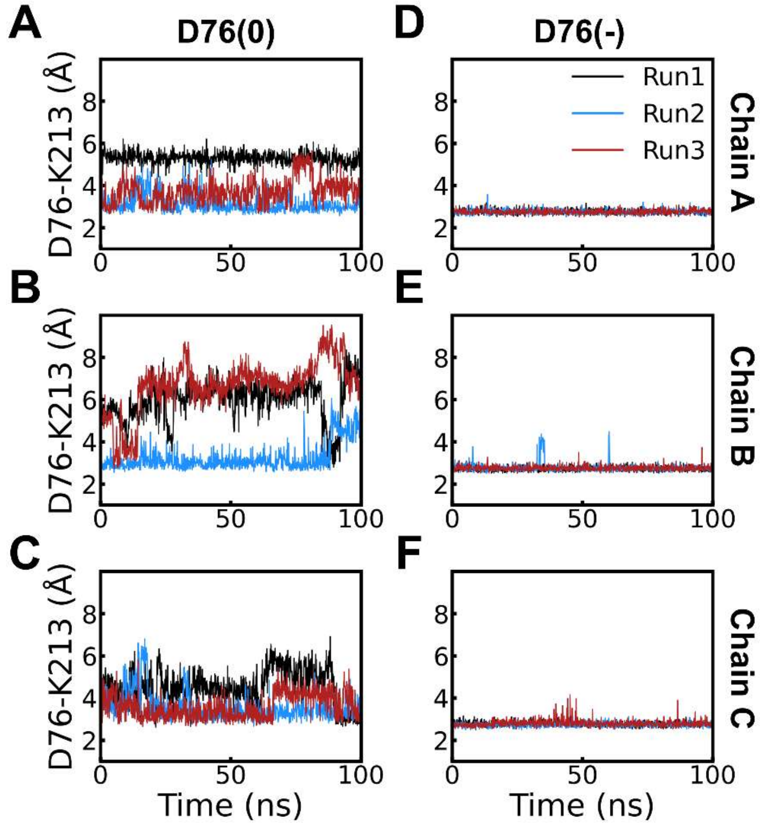


**Figure S6.** MD simulation trajectories of the minimal distance between Asp76 and Lys213 side chains in the proton pump NsXeR where Asp76 is fixed in (A-C) protonated and (D-F) deprotonated states, respectively. Simulations started from the crystal structure (PDB code: 6EYU).

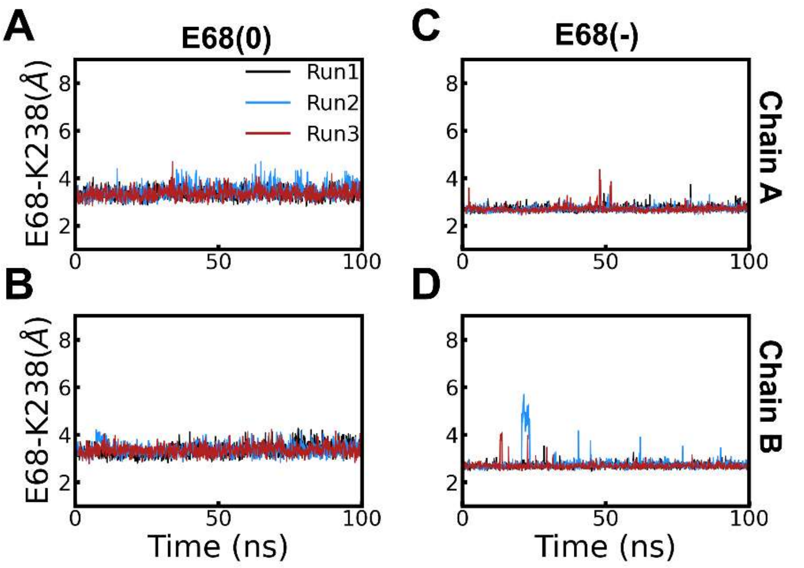


**Figure S7.** MD simulation trajectories of the minimal distance between Glu68 and Lys238 side chains in the anion channel iC++ where Glu68 is fixed in (A-B) protonated and (C-D) deprotonated states, respectively. Simulations started from the crystal structure (PDB code: 6CSM).

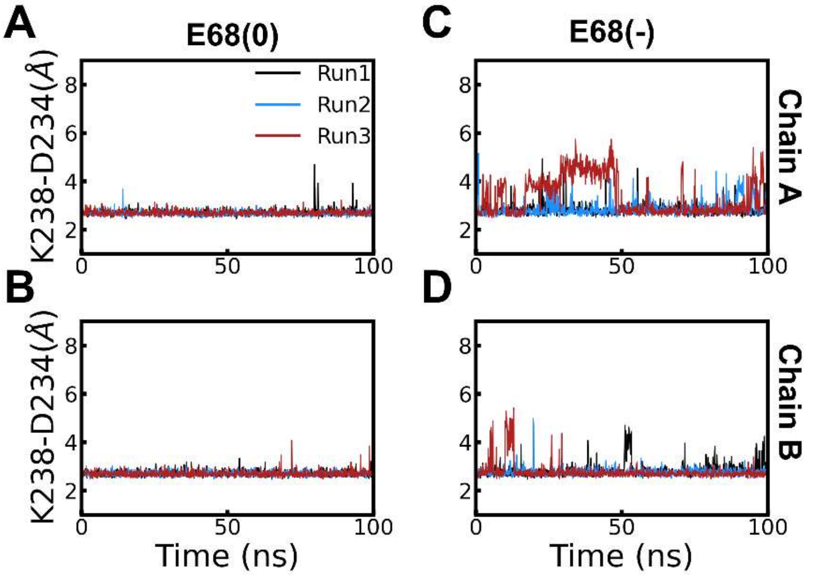


**Figure S8**. MD simulation trajectories of the minimal distance between Asp234 and Lys238 side chains in the anion channel iC++ where Glu68 is fixed in (A-B) protonated and (C-D) deprotonated states, respectively. Simulations started from the crystal structure (PDB code: 6CSM).

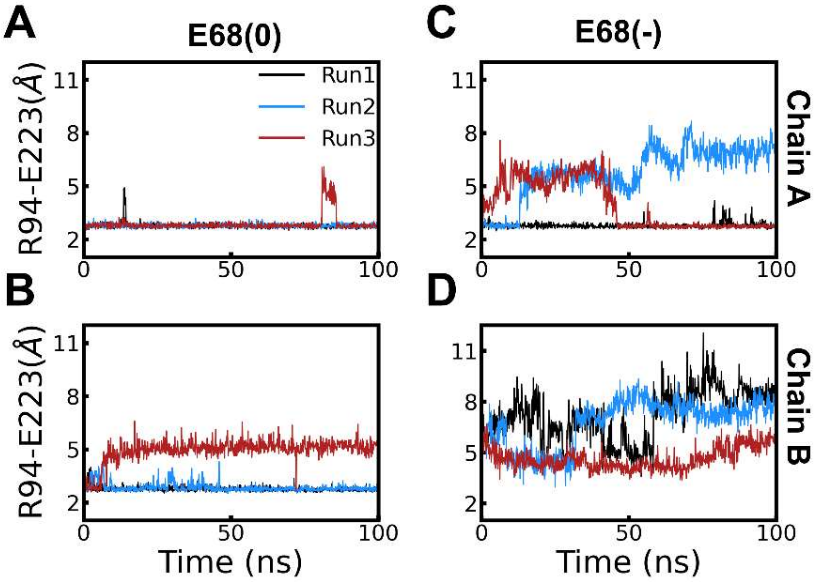


**Figure S9.** MD simulation trajectories of the minimal distance between Glu223 and Arg94 side chains in the anion channel iC++ where Glu68 is fixed in (A-B) protonated and (C-D) deprotonated states, respectively. Simulations started from the crystal structure (PDB code: 6CSM).

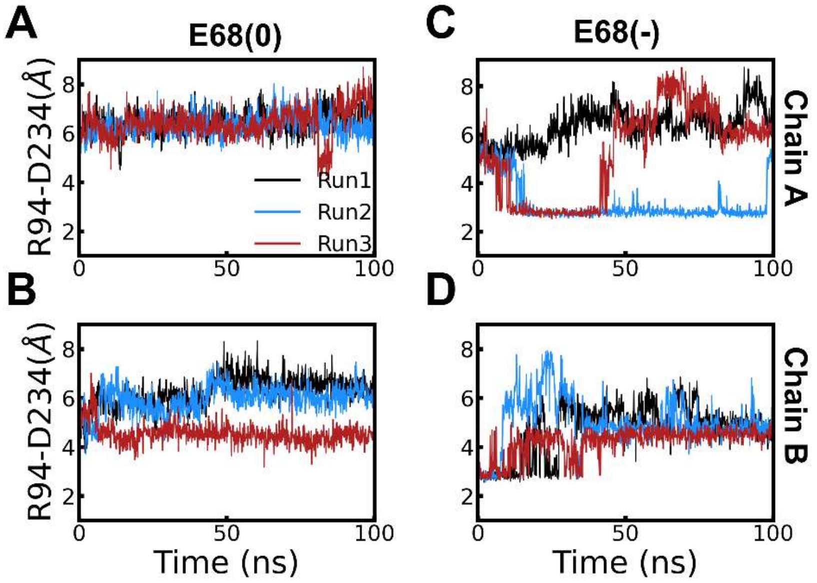


**Figure S10.** MD simulation trajectories of the minimal distance between Asp234 and Arg94 side chains in the anion channel iC++ where Glu68 is fixed in (A-B) protonated and (C-D) deprotonated states, respectively. Simulations started from the crystal structure (PDB code: 6CSM).

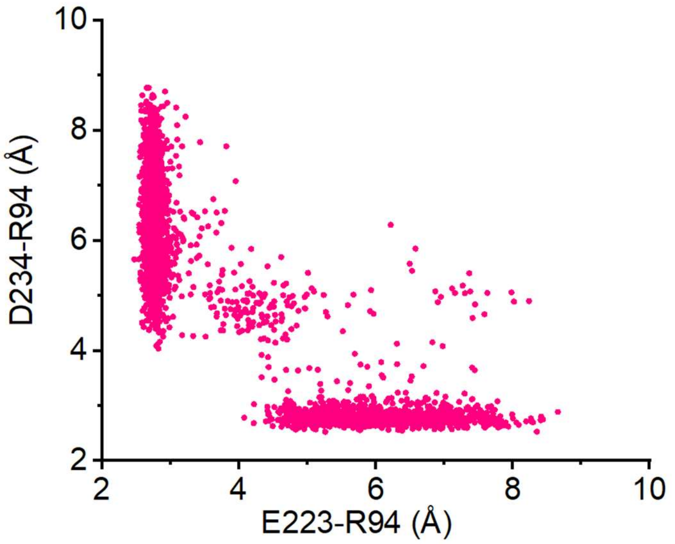


**Figure S11.** Correlation plot of minimal distances between Glu223 and Arg94 side chains (E223-R94) and between Asp234 and Arg94 side chains (D234-R94). Data come from Fig. S9C and Fig. S10C above.

### Molecular Dynamics (MD) Simulation

#### System Preparation

The Membrane Builder module of the CHARMM-GUI interface was invoked to build up the simulation system[6-8]. Truncated N- and C-terminals were acetylated and amidated with $CH_3CO$ (ACE) and $NH_2$ (CT2), respectively. Protonation states of Asp, Glu, His, and Lys residues under the physiological pH were assigned based on p$K_a$'s extracted from MPKaDB. The simulated pH for $GlcP_{Se}$, NsXeR and iC++ are 5.5[9], 7.0[10] and 6.5[11] , respectively. Given pH, protonation states for Asp, Glu, His and Lys were determined by p$K_a$'s in MPKaDB, except for Asp22 in GlcP, Asp76 in NsXeR and Glu68 in iC++ that were selected to study the possible influences of their charge states on salt bridge networks. As to Tyr, Cys and Arg, reference p$K_a$'s were utilized. The protein was embedded in a lipid bilayer composed by 1-palmitoyl-2-oleoyl-*sn*-glycero-3-phosphocholine (POPC) lipids.The default rectangular box type was selected, allowing the box size to adjust at membrane normal or z axis. Then box length at z axis was estimated according to the default water thickness of 22.5 Å capped to both sides of the protein in z direction. Apart from the bulk solvent, water molecules were added to fill cavities if any in pores. Finally, sodium and chloride ions were added to neutralize the simulation system and simultaneously mimic the physiological ionic strength of 0.15 M.

#### Simulation Protocols

CHARMM36m[12] and CHARMM36[13] force fields were selected to describe proteins and lipids, respectively. CHARMM-modified TIP3P model was utilized to mimic water molecules[14,15]. Given periodic boundary conditions, the particle mesh Ewald algorithm was utilized to describe electrostatic interaction with a real-space cutoff of 12 Å[16]. As to the reciprocal space, the 1-Å grid spacing and sixth-order spline interpolation were considered. The force switch scheme was employed to deal with the van der Waals interaction where the switching on and off distances are 10 and 12 Å, respectively. Before production runs, the equilibration protocol recommended by CHARMM-GUI was employed to relax the simulation systems, particularly lipids. Given a protonation state of the screened Asp, three independent simulations that lasted 100 nanoseconds were carried out for each protein, which leads to an amount of 18 simulations. Simulations were performed under the NPT ensemble using the OpenMM engine[17]. Specifically, the Hoover thermostat was employed to maintain a constant temperature of 310 Kelvin[18]. For pressure control, the Langevin piston coupling method was applied to maintaining the system at 1 atm[19]. To allow a time step of 2 fs, the SHAKE algorithm was adopted[20].
